# Cultural Evolution of Genetic Heritability

**DOI:** 10.1101/2020.06.23.167676

**Authors:** Ryutaro Uchiyama, Rachel Spicer, Michael Muthukrishna

## Abstract

Behavioral genetics and cultural evolution have both revolutionized our understanding of human behavior, but largely independently of each other. Here we reconcile these two fields using a dual inheritance approach, which offers a more nuanced understanding of the interaction between genes and culture, and a resolution to several long-standing puzzles. For example, by neglecting how human environments are extensively shaped by cultural dynamics, behavioral genetic approaches systematically inflate heritability estimates and thereby overestimate the genetic basis of human behavior. A WEIRD (Western, educated, industrialized, rich, democratic) gene problem obscures this inflation. Considering both genetic and cultural evolutionary forces, heritability scores become less a property of a trait and more a moving target that responds to cultural and social changes. Ignoring cultural evolutionary forces leads to an over-simplified model of gene-to-phenotype causality. When cumulative culture functionally overlaps with genes, genetic effects become masked, or even reversed, and the causal effect of an identified gene is confounded with features of the cultural environment, specific to a particular society at a particular time. This framework helps explain why it is easier to discover genes for deficiencies than genes for abilities. With this framework, we predict the ways in which heritability should differ between societies, between socioeconomic levels within some societies but not others, and over the life course. An integrated cultural evolutionary behavioral genetics cuts through the nature–nurture debate and elucidates controversial topics such as general intelligence.

## 1. INTRODUCTION

Business is booming in behavioral genetics. We’re in the midst of a genome-wide association gold rush (Visscher et al., 2017). The availability of powerful computers and sequenced DNA of millions of people has led to a frenzied search for single nucleotide polymorphisms (SNPs) that correlate with a variety of psychological and behavioral traits (Horwitz et al. 2019; Harden and Koellinger 2020; Mills and Tropf 2020). These range from memory capacity (Papassotiropoulos et al. 2011), cognitive ability (Coleman et al. 2019) and educational attainment (Lee et al. 2018) to moral attitudes (Brandt and Wetherell 2012), political orientation (Hatemi et al. 2011), temporal discounting (Sanchez-Roige et al. 2018), socio-economic status (Hill et al. 2016), temperament (Zwir et al. 2018), and happiness (Wingo et al. 2017). These correlations are highly statistically significant (typically at least *p* < 5 ×10^−8^; Fadista et al., 2016) and the curse of reverse causality has apparently been lifted. As Plomin and von Stumm (2018) put it, genome-wide polygenic scores “are an exception to the rule that correlations do not imply causation in the sense that there can be no backward causation…nothing in our brains, behavior or environment changes inherited differences in our DNA sequence.”

The last two decades have also seen a parallel revolution in cultural psychology and cultural evolution that have identified cultural correlates of our psychology and behavior (Henrich, Heine, and Norenzayan 2010; Nisbett 2003; Muthukrishna et al. 2020; Henrich 2016; Muthukrishna and Henrich 2019; Gelfand 2018). These range from fairness and prosocial norms (Henrich, Heine, and Norenzayan 2010; Schulz et al. 2019) and attribution of blame (Barrett et al. 2016) to perceptual style (Kitayama et al. 2003), susceptibility to visual illusions (Henrich, Heine, and Norenzayan 2010), numeric chunking (Domahs et al. 2010), interpretation of linear and logarithmic numeric scales (Dehaene et al. 2008), neural correlates of reading (Bolger, Perfetti, and Schneider 2005; Tan et al. 2005), event segmentation (Swallow and Wang 2019), working memory (Amici et al. 2019; Guida et al. 2018), spatial cognition (Majid et al. 2004), motor development (Karasik et al. 2015), folkbiology (Medin and Atran 2004; Waxman, Medin, and Ross 2007), and personality (Smaldino et al. 2019; Gurven et al. 2013). Cultural evolution is part of a broader theoretical framework—dual inheritance theory ^1^ — that incorporates genes, environment, culture and learning to explain and predict human psychology and behavior (Muthukrishna and Henrich 2019). This body of research suggests that humans not only have a genetic inheritance from their parents, as all animals on Earth do, but also a cultural inheritance from their societies. Genes, culture, and the environment have often co-evolved, shaping many aspects of human psychology and behavior (Henrich 2016; Laland 2018).

These revolutions in behavioral genetics and cultural evolution have largely occurred independently of each other. Cultural evolution has incorporated some aspects of behavioral genetics (e.g. Laland, Odling-Smee, and Myles 2010; Creanza, Kolodny, and Feldman 2017; Feldman and Ramachandran 2018), but not to any significant degree. Behavioral genetics in turn has been largely agnostic with respect to cultural evolution. Nonetheless, cultural evolution has implications for the interpretation of findings in behavioral genetics. Specifically, cultural evolution suggests that behavioral genetics overestimates the predictive power of genes. This overestimation is obscured by a WEIRD gene problem that severely restricts the range of sampled environments. In this article, we reconcile cultural evolution with behavioral genetics. This reconciliation offers new interpretations for various puzzles, such as differences in heritability between and within populations, differences in heritability across development, and general intelligence. In doing so, it challenges the current interpretation of the roles of genes, environment, and culture in behavior.

## 2. PROBLEMS INTERPRETING GENES, ENVIRONMENT, AND CULTURE

### 2.1. Causal locus problem

Natural selection selects for genes that confer a fitness advantage in an environment. Genes, therefore, are for something. That is, genes have a function or functions to be discovered and mapped (e.g. The Gene Ontology Consortium 2017). Genes, of course, interact with the environment, but the “gene for” intuition permeates how we talk about genes, how we represent them in diagrams, and how we measure their effects. In cultural evolution, this genetic primacy gives way to explicit formalizations of essential interactions between genes, culture and the environment.

Culture and genes are interwoven in the construction of many behavioral traits, making separation impossible. Is language primarily the result of culture or genes (Dediu 2011; Christiansen and Chater 2008; Chater, Reali, and Christiansen 2009)? This kind of functional entanglement occurs across various levels of biological organization: between genes within the same genome— *intragenomic* (Phillips 2008), between nuclear and organellar (mitochondria and plastid) genomes—*cytonuclear* (Sloan et al. 2018), and between host and microbial symbiont genomes— *holobiontic* (Bordenstein & Theis, 2015). Mitochondria, for example, are believed to have undergone extensive reductive evolution, transferring nearly all of their genes to the nuclear genome (Wolf and Koonin 2013; Sloan et al. 2018). Indeed, the residual mitochondrial and nuclear genomes collaboratively assemble “chimeric” proteins (Osada and Akashi 2012). This kind of coevolution is common, undermining the tenets of the Modern Synthesis that enshrine the causal supremacy of nuclear DNA (Laland et al. 2015; Jablonka and Lamb 2005). Functional interweaving also occurs beyond genes alone, between genes and the environment.

One clear example is the *GLO* gene, which is required to endogenously synthesize vitamin C. Vitamin C is an essential nutrient and the *GLO* gene is present across most of the animal kingdom. However, vitamin C synthesis is metabolically costly, and *GLO* is inactive in some species that have access to sufficient quantities of the nutrient in their diets (Drouin, Godin, and Page 2011). These include taxa such as teleost fishes, guinea pigs, many bats, some passerine birds and anthropoid primates, i.e. monkeys and apes (Chatterjee 1973). Anthropoids for instance occupy a frugivorous niche, and fruits often contain sufficient vitamin C. Here gene function is offloaded onto environmental resources. In turn, this offloading has behavioral implications. If a species becomes dependent on its environment (auxotrophic) for vitamin C, both its behavioral range and evolutionary trajectory become constrained by the availability of the nutrient. Humans are a nice example of this. As our species migrated across the planet, we found ourselves in environments where vitamin C was in short supply. A deficiency of vitamin C causes scurvy—the bane of seafarers until the trial-and-error discovery that certain food items like sauerkraut and citrus could prevent ships from being packed with tired, bleeding, toothless, and eventually dead sailors (Lamb, May, and Harrison 2017).

When a single change in genes, culture, or environment creates a problem (as is the case of *GLO*, vitamin C, and scurvy), that gene, culture, or aspect of the environment appears to be causally responsible. But phenotypic traits are the result of complex interactions between and within these domains. This creates a ‘causal locus’ problem that obscures the complex causal basis of a trait and instead presents the appearance of a singular cause, because a single change can break a system, but many factors contribute to making it.

#### 2.1.1 Genes that break and genes that make

The more complex a system, the more ways it can fail. Take the history of lighting: compared to the two ways in which a wood-fueled fire can be extinguished (smothering and exhaustion of fuel), there are 7 known failure modes for a fluorescent bulb and more than 30 for the newer LED bulb (de Groot et al. 2013). A faulty rubber O-ring caused the space shuttle Challenger to explode, and a severed fiber-optic cable knocked out internet access for a large swath of people across India and the Middle East. There is a fundamental asymmetry between the identification of elements that support a system and those that undermine it. A well-functioning system is the product of a design process that has solved many problems and closed many paths that do not work. It is therefore easier to find ways to break the system than ways to build it.

The same is true for genes that build life. Organisms are the outcomes of complex, emergent interactions involving many genes and their surroundings (Davies 2014), but there are many ways these interactions can go wrong. It is easier to identify deleterious genetic mutations than beneficial mutations, as deleterious mutations are more common and have larger effects—the space of failure is larger than the space of success. For example, identifying the cause of Mendelian disorders such as cystic fibrosis and Huntington’s disease is relatively simple, as they are caused by mutations in a single gene. But it is much harder to identify genes that are responsible for constructing a complex trait. For instance, there is no single gene for intelligence; recent analyses with samples sizes in the hundreds of thousands have detected >1000 genes linked to intelligence (Savage et al. 2018; Davies et al. 2018), with the actual number likely to be much higher. Each gene only explains a miniscule fraction of variation in intelligence, and is unlikely to contribute in a clear causal path. Indeed, we have no systematic understanding of exactly how these genes create intelligence. We are unlikely to develop this understanding through the main effects of regressions or even higher-order statistical interactions, although such techniques can identify genetic correlates of intelligence. In contrast, the causal mechanisms behind single gene mutations that cause intellectual disability— e.g. *BCL11A* (Dias et al. 2016), *PHF8* (Bathelt et al. 2016), *ZDHHC9* (Schirwani et al. 2018)— are relatively well understood.

A gene can be beneficial in one environment but not in another. For example, we have known for a long time that increasing nutrition (Lynn 1990; Stoch et al. 1982), improving schooling (Ceci 1991; Davis 2014; Ritchie and Tucker-Drob 2018), and removing parasites (Wieringa et al. 2011) have positive effects on general intelligence. None of this is surprising, but it means that in a society where parasite infection is kept under control, we would not notice that parasite status correlates with intelligence, due to a lack of sufficient variation in parasite load. For the same reason, a correlation between lead exposure and IQ (Needleman and Gatsonis 1990; Wasserman et al. 1997) will not be revealed in a society where lead is not a problem. The same principle applies to genes: genes that provide protection against malnutrition, parasites, or pollution would only be positively associated with intelligence in environments where these insults occur. In environments where these challenges have been overcome, the same genes would not be associated with intelligence, and can even be deleterious. For example, being a carrier (heterozygous) for abnormal hemoglobin via sickle cell trait (Elguero et al. 2015) or thalassemia (Mockenhaupt et al. 2004) protects against malaria and is thus beneficial in an environment with the *Plasmodium falciparum* parasite. Because malaria is known to have a negative impact on cognitive development (Holding and Snow 2001), we would expect the gene for abnormal hemoglobin to be positively associated with intelligence in environments with a high risk of malaria. As the risk of malaria decreases heterozygosity will be neutral or deleterious, but this too depends on environmental factors such as diet. Similarly, alleles that protect against parasite infection (Carter 2013) or lead poisoning (Onalaja and Claudio 2000) will be predictive of IQ only if the environmental risk factors are present in sufficient quantities. In an environment with arsenic contamination, variants in *AS3MT* associated with more efficient arsenic metabolism (Schlebusch et al. 2015) may be predictive of intelligence (Wang et al. 2007).

In many of these examples, culture masks genetic effects. The statistical association between gene and trait is dependent upon environmental context. The more protective factors are accumulated within a culture, the more masking there will be and the easier it will be to identify harmful environmental factors that can undermine a trait compared to constructive factors that can boost it; a protective environment is built to do more work than a deprived environment and can therefore fail in more ways. Only when we peel back the comforts enjoyed by advanced economies, will genes that confer advantage under conditions like malnutrition or disease threat become unmasked and detectable. Because such environmental deprivation undermines performance on IQ tests, these genes will be identified by behavioral geneticists as genes for cognitive ability.

For most of human history, what we now call “deprived” conditions were the norm. Two hundred years ago, 89% of humanity lived in extreme poverty (Ravallion 2016), 88% were illiterate (van Zanden et al. 2014), and 43% of children died before they were five years old (Gapminder 2020). Conditions have rapidly improved: rates of extreme poverty are now 10%, illiteracy is down to 14%, and deaths before five years of age are now 4% (‘World Bank Group - International Development, Poverty, & Sustainability’ 2020; UNESCO Institute for Statistics 2013). Our world still suffers from immense global inequality even if most are now better off. When a genome wide association study (GWAS) identifies genes for intelligence, prosociality, or longevity, those genes should not be assumed to be valid beyond a specific people, living in a specific environment, in a specific time. This explains why candidate genes often do not replicate outside the population in which they are discovered (Samek et al. 2016; Dumas-Mallet et al. 2016; Duncan and Keller 2011; Culverhouse et al. 2018).

This argument goes beyond simply a mismatch between genes and culture. Instead, genes can only be evaluated with respect to the cultural environment in which they are expressed. For this reason, heritability is less a property of a trait, and more a combined measure of features of the environment including culture, their respective variances, and the causal relationship between genes and traits.

#### 2.1.2 Interpreting heritability

To illustrate the challenges in understanding how genes, environment and culture affect heritability, we begin with a relatively simple example: the heritability of cancers associated with skin pigmentation. Genes affect the level of skin pigmentation and propensity for tanning instead of burning. These are ancestral adaptations to levels of UV radiation at different latitudes (Barsh 2003; Sturm and Duffy 2012). Darker pigmentation protects against high levels of UV radiation, such as near the equator. Lighter pigmentation enables vitamin D synthesis in low levels of UV radiation, such as at Northern latitudes (Jablonski and Chaplin 2010; 2017). It is important to get the correct amount of UV radiation—too much causes skin cancer, but too little causes vitamin D deficiency, which is associated with other cancers (Garland et al. 2006; Edlich et al. 2009).

Worldwide migration has led to people with skin pigmentation mismatched to the level of UV radiation: Australians with European ancestry have higher rates of skin cancer than Australian Aboriginals and other non-white populations (Australian Institute of Health and Welfare 2016), and conversely, Europeans with African and South Asian ancestry have higher rates of vitamin D deficiency and associated cancers (Cashman et al. 2016; Spiro and Buttriss 2014). However, cultural practices and technologies can provide non-genetic adaptations to these mismatches: fairer Australians wearing sunscreen, a hat and covered clothing (Montague, Borland, and Sinclair 2001)^2^ and darker Europeans consuming vitamin D supplements and vitamin D-rich or fortified foods.

In this example, the challenges to interpreting heritability and understanding GWAS results are perhaps more obvious than in many psychological cases. The heritability of skin cancer should be highest when there is more diversity of skin pigments, more homogeneity of cultural practices, and high UV radiation. For example in the US, the heritability of Vitamin D concentrations is approximately 70% during winter, but effectively 0% during summer (Karohl et al. 2010). Heritability is not only a function of the causal paths between genes and traits, but also a function of the cultural environment and its variability. GWAS also suffer from this problem when correlating SNPs with traits—SNPs for skin pigmentation are identified more easily where there is diversity in skin pigmentation. We illustrate this example in Figure 1 below.

**Figure 1:**
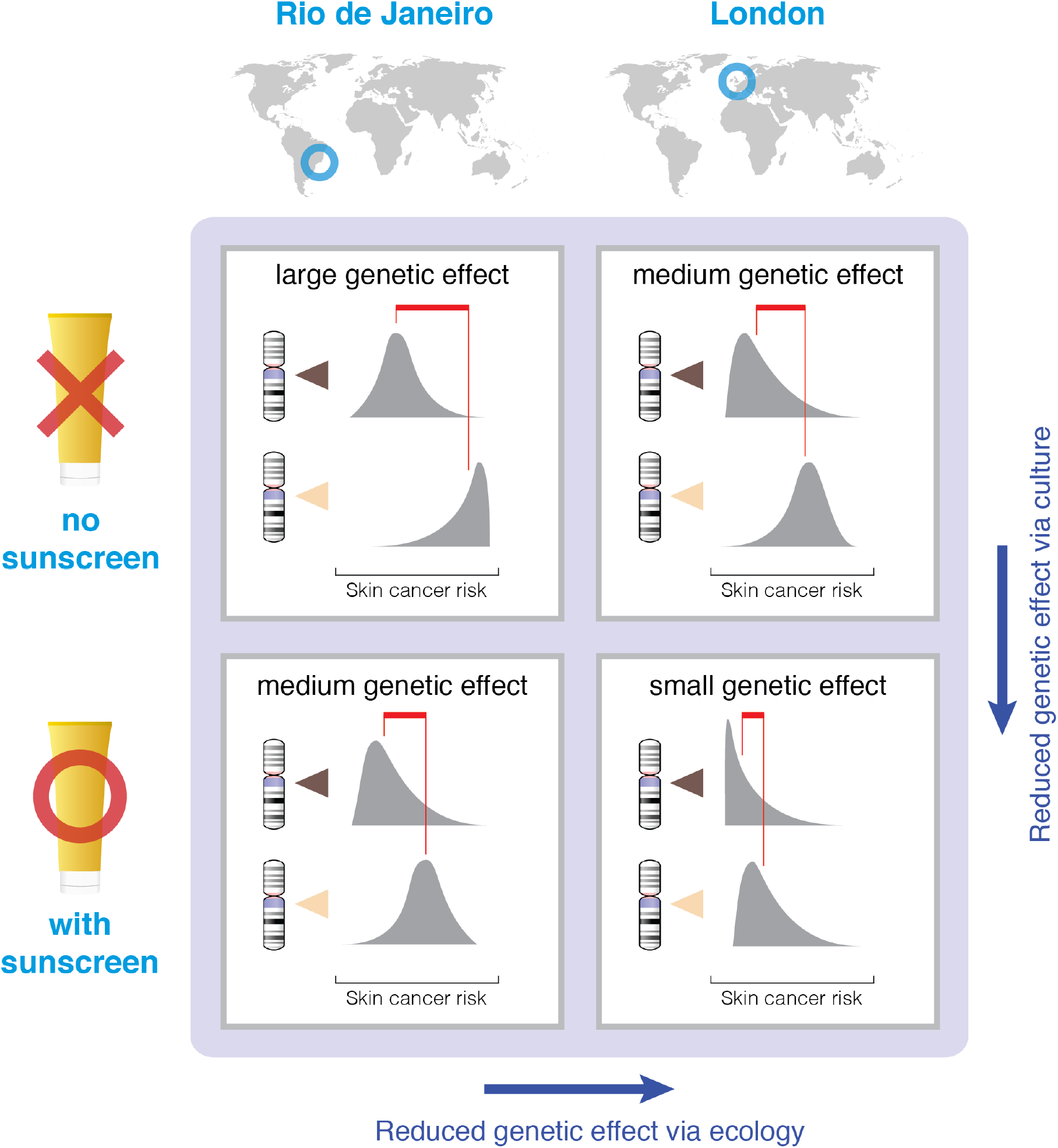
An illustration of the effect of sunscreen and geographic location on the effect size of a skin pigmentation gene with respect to skin cancer risk. The largest genetic effect should be found in societies that lack sunscreen and reside in locations with high levels of UV radiation (top-left square). Genetic effects should be reduced with either the introduction of sunscreen or residence in a lower-UV environment, both factors that mask the effect of skin pigmentation (bottom-left and top-right squares, respectively). The smallest effect should be found in societies that have both low UV and sunscreen (bottom-right square). Chromosomes with dark indicators represent genes for strong pigmentation, and those with light indicators represent genes for light pigmentation. Gray distribution represents population distributions for skin cancer risk, and red lines point to the mean of each distribution.

That heritability is affected by the environment is obvious and well understood (Feldman and Ramachandran 2018; Hamer and Sirota 2000; Moore and Shenk 2016; Turkheimer, Pettersson, and Horn 2014; Vitzthum 2003; Tenesa and Haley 2013; Charmantier and Garant 2005). Lewontin and colleagues (Lewontin 1970a; 1974; Feldman and Lewontin 1975) long ago described the fallacy of extrapolating heritability scores from one population to another. Their argument was made from the standpoint of gene-environment interactions: genetic effects must be understood in the environmental conditions under which the genes are expressed. What is less well understood is the systematic way in which the environment changes over time, and how this affects the interpretation of heritability—that is, the cultural evolution of genetic heritability (Figure 2).

**Figure 2:**
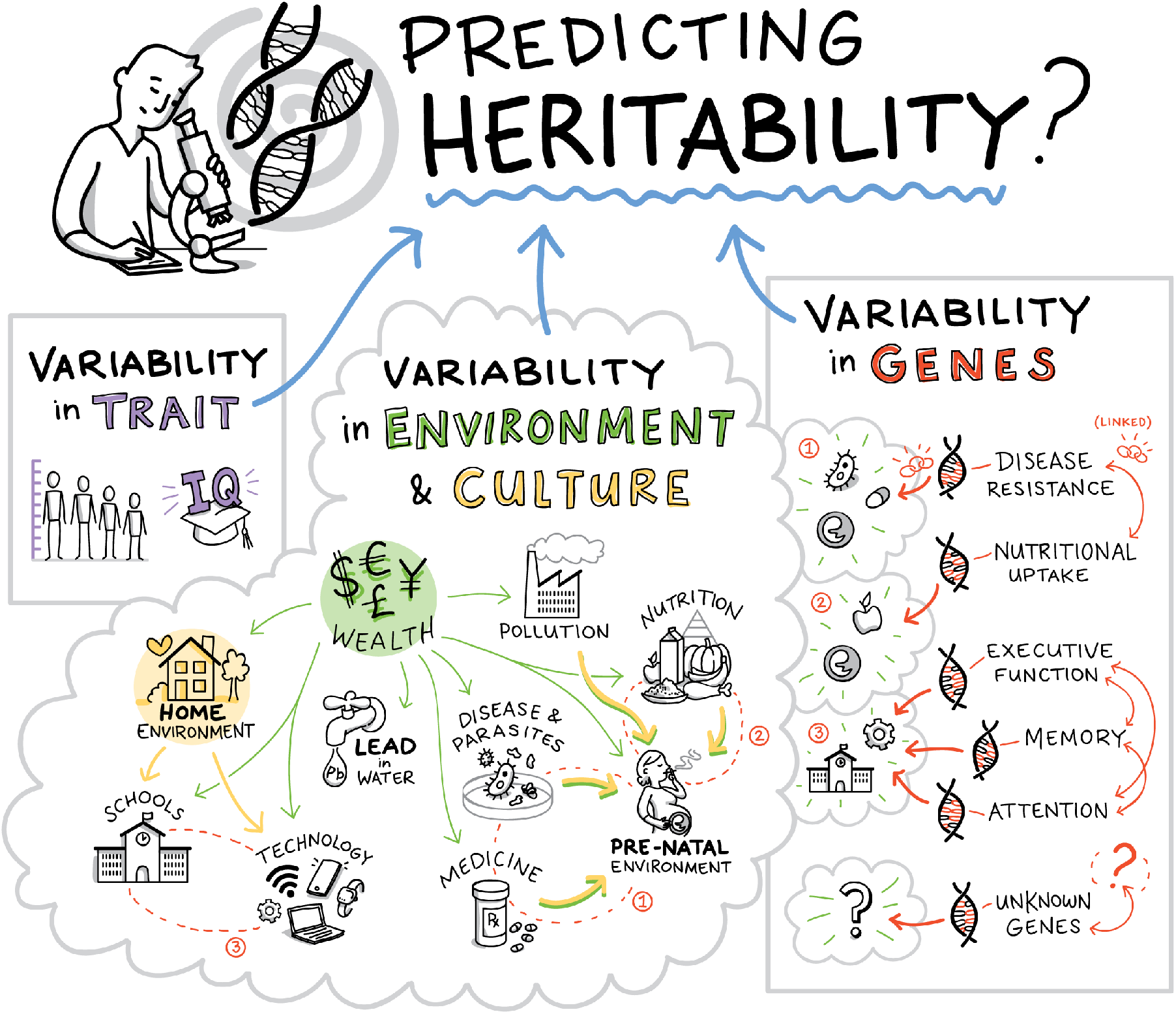
Genetic heritability is a function of variability in the phenotypic trait, variability in the environment, including the cultural environment, and variability in genes. Although heritability is commonly interpreted as a genetic effect, cultural evolution and diffusion can also impact the variability of environmental variables, and thus heritability. Psychological and behavioral phenotypes are typically the outcome of a complex network of interactions that involve all these factors.

The example of skin pigmentation and cancer should make it clear that there is no overarching, one-quantity heritability of a trait to be discovered. There is no fixed answer to the question, “What is the heritability of skin cancer?” We can attempt to control for latitude (environment) or annual sunscreen sales (culture), but to do so is to privilege genes where there is little reason to do so. That is, there is no sensible or useful ‘raw’ heritability that is independent of culture. The effect of culture is many-fold, from the amount of time we spend indoors doing clerical work or outdoors on our daily running route, to the kind of clothes we wear and the amount of antioxidants in our diet (Katta and Brown 2015; Ichihashi et al. 2000; Godic et al. 2014). The question, “Which SNPs are associated with skin cancer?” is similarly culturally dependent. In societies where sunscreen use is common, we expect genes that govern skin pigmentation to be less predictive of skin cancer compared to societies where it is not. We expect genes that control antioxidant metabolism (Oskina et al. 2014) to be less predictive of skin cancer in societies whose foods are rich in antioxidants—such as in traditional Mediterranean cuisine (Visioli and Galli 2001). The presence of a cultural trait can mask or amplify a genetic effect.

If culture acts upon environmental factors in such a way that results in a reduction of the phenotypic variation explained by a particular SNP, then this SNP–phenotype association will be diminished, and the other SNPs that contribute to that same phenotype will be proportionately inflated. Therefore, sunscreen would be expected to inflate the perceived contribution of genes that control antioxidant metabolism, even if UV has historically been the more significant cause of skin cancer. Such cultural effects are, however, not limited to specific traits and specific genes. Cumulative cultural evolution is responsible for genetic inflation across all observable human environments, and thereby biases the assessment of genetic effects across the board.

### 2.2. Cultural compression problem

The cultural compression problem identifies a basic selection bias that emerges as a byproduct of cumulative cultural evolution. The problem is not simply that gene–environment interactions complicate our understanding of genetic effects, it is that for humans, there is no ‘baseline’ environment against which true genetic effects can be neutrally evaluated in the first place.

Assume that for a given society we were able to collect comprehensive data on genetic effects across all relevant environmental variables that contribute to some trait. This would allow us to exhaustively map out the ‘reaction norms’ that specify expected phenotypic outcomes over the full range of extant genetic and environmental variation. But even this complete set of observable gene– environment interactions would not fulfil the role of a neutral backdrop for the assessment of true genetic effects. This is because our measurements do not map out the dimensions of an ‘adaptive landscape’ (Simpson 1944; Wright 1932) as we might imagine; instead they map out the local and global peaks that have already been climbed by cumulative cultural evolution.

Gene-environment interactions are not sufficient, as any environment that we observe in a human sample is always the post hoc outcome of a particular, idiosyncratic cultural history. Given the open-ended and path-dependent nature of the evolutionary process (Taylor et al. 2016; de Vladar, Santos, and Szathmáry 2017; Coffman 2014; Gould 1989), combined with the huge scope for variation in cultural traits compared to genetic traits (Bell, Richerson, and McElreath 2009; Muthukrishna et al. 2020), we expect that observations of environmental variables in extant human populations will represent only a small subset of possible, viable environments (Figure 3).

**Figure 3:**
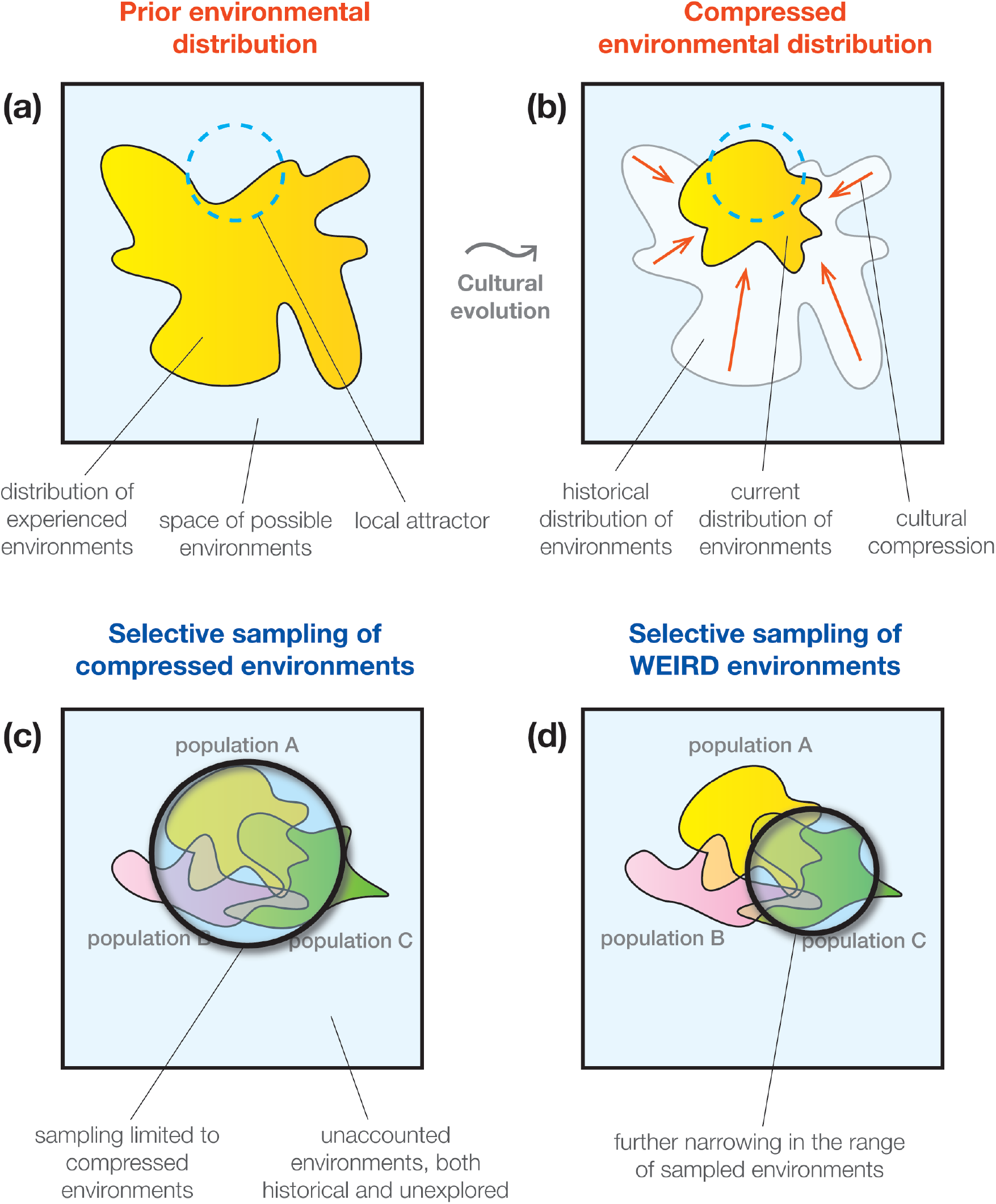
An illustration of the cultural compression problem. (a) The yellow region represents the distribution of experienced environments of a hypothetical society at a past time point. The unoccupied light blue area represents the unexplored regions of the space of possible, viable environments. The blue ring represents a set of environmental states that are better adapted to ecological challenges and functions as a local attractor on an adaptive landscape. (b) The environmental distribution at a later time point. Through cultural evolutionary dynamics such as conformist transmission and selective imitation, the society has converged around the local attractor. (c) Even if researchers were able to obtain samples from all extant populations, their observations would be limited to a particular subspace of possible environments that is contingent upon cultural history. Because genetic effects can only be evaluated with respect to particular environments, genes may have vastly different effect sizes or functions outside of this observable range. (d) In practice, researchers conduct the majority of their analyses within a handful of societies that represent a small fraction of global genetic and cultural variation. This limitation further narrows down the range of observed environments and thus impedes generalizability of genetic effects.

The enormous capacity that we have to shape our own environments through cultural transmission is a defining characteristic of our species (Laland and O’Brien 2011; Boyd, Richerson, and Henrich 2011). When we condition our observations on extant environments, we subject ourselves to a selection bias whose severity reflects the magnitude of impact that cultural evolution has had upon our environments. Because observations can only occur after the iterated cultural diffusion of traits, environments are necessarily more homogenous than would be expected in the absence of culture, and heritability estimates are necessarily inflated when using standard behavioral genetic methods. It is not particular cultural inventions that inflate heritability, but rather the process of cumulative cultural evolution itself.

As an illustration of how cultural transmission inflates heritability, consider the effect of standardized education. When Samuelsson et al. (2008) measured the heritability of reading and spelling test scores, Australian twins demonstrated a narrow-sense heritability of 0.84 in kindergarten and a similar score of 0.80 in Grade 1. But in their cohort of Swedish and Norwegian twins, heritability was only 0.33 in kindergarten, but rose to 0.79 in Grade 1. Heritability was at the same level in both the Australian and Scandinavian children in Grade 1, but not in kindergarten. Why? Australian children begin receiving compulsory literacy instruction in kindergarten, while in Scandinavia the kindergarten curriculum emphasizes social, emotional, and aesthetic development—literacy instruction only begins in Grade 1. Therefore, Australian kindergarteners are exposed to standardized environmental input and much of the remaining variation in reading ability is explained by genetic differences, whereas for the Swedish and Norwegian kindergarteners, variation in the amount of reading instruction received at home is much larger than any genetic differences. In line with this interpretation, Samuelsson et al. (2008) show that the boost in heritability among the Scandinavian children was also accompanied by an almost equivalent decrease in phenotypic variance attributed to the common (home) environment, which would include home instruction.

These results demonstrate how heritability does not depend on the intrinsic properties of genes alone. Heritability measures the effect of genes against a background of environmental variation: it is highest when the relevant environmental input is uniform across a sample, and shrinks as environmental input becomes more varied. In the present example, it is standardized education that flattens environmental variance, but any mechanism that contributes to the homogenization of relevant environmental features will amplify heritability. When we say that the heritability of reading among Scandinavian children jumps up to 0.79 when they enter first-grade, this measurement reveals just as much about the power of modern schooling as it does about the genetic basis of literacy.

Compulsory education is a particularly powerful channel of enculturation, as it delivers a standardized package of skills and knowledge across virtually all individuals in an age-cohort. But if we were to assess the genetic basis of literacy skill in schoolchildren without accounting for the impact of their particular educational curricula on environmental variation, we would be subjecting ourselves to a selection bias, with no idea of what the magnitude of this bias might be. This would distort our understanding of the generalizability of our finding to samples that have undergone different educational curricula, and even more so to those with different levels of educational attainment. Note that even the literacy instruction provided in the home environment is already shaped by cultural evolution, both in terms of the content being transmitted (reading and spelling), and the structures that are transmitting (family organization in Western countries; Schulz et al. 2019; Henrich 2020). In societies that produce literate children, culture impacts the heritability of literacy from the moment that variation in this trait emerges in development, virtually sealing off the possibility of assessing ‘baseline’ heritability even at the very start of life. As such, the cultural compression problem is compounded by limited sampling of both genes and environments.

### 2.3. WEIRD gene problem

Behavioral genetics suffers from its own variant of the Western, educated, industrialized, rich, democratic (WEIRD) people problem, which was originally raised in the field of experimental psychology (Henrich, Heine, and Norenzayan 2010). The WEIRD people problem refers to the vast over-representation in published studies of individuals from developed Western countries, who are all similar in their cultural history, social values, and standards of living^3^. Behavioral genetic samples are both culturally WEIRD and genetically WEIRD.

A comprehensive meta-analysis that claims to contain essentially all twin studies published between 1958 to 2012 (Polderman et al. 2015) reveals that 94% of sampled twin pairs were from Western populations. The United States, United Kingdom, and Australia alone accounted for almost 60%, and Nordic countries accounted for another 25%. Of the non-Western countries (6%), two thirds (4%) are from northeast Asia—specifically, China, Japan, South Korea, and Taiwan, countries that are not Western, but have most of the remaining letters of the WEIRD acronym. The remainder of the world, representing the vast majority of the human population, accounts for only 2% of the dataset.

GWAS too suffers from a WEIRD gene problem (Need and Goldstein 2009; Popejoy and Fullerton 2016; Sirugo, Williams, and Tishkoff 2019). As of 2017, 88% of samples in GWAS were of European ancestry (Mills and Rahal 2019)^4^. Paralleling the twin studies data, 72% of participants were recruited from just three countries—US, UK and Iceland—with nearly 20% of the remainder being recruited from Japan, China and South Korea.

Genetic samples largely come from WEIRD countries that are clustered together along multiple cultural dimensions (Hofstede 2001; Inglehart and Welzel 2005; Muthukrishna et al. 2020), and that are perhaps an extreme unrepresentative outlier on many psychological and behavioral measures, with these countries registering the highest scores for traits like individualism, analytical thinking, and prosociality toward strangers, and the lowest scores on opposite constructs such as collectivism, holistic thinking, and prosociality toward relatives but not strangers (Henrich, Heine, and Norenzayan 2010; Muthukrishna et al. 2020; Schulz et al. 2019; Henrich 2020). When we restrict the scope of genetic samples, the cultural environment against which genetic effects are evaluated also become skewed. This greatly reduces the interpretability of genetic effects, as cumulative culture obscures the causal locus of phenotypic outcomes.

This WEIRD gene sampling bias in GWAS is deeply problematic. Polygenic scores do not translate well across ancestry groups (Bitarello and Mathieson 2020; Guo et al. 2019; Curtis 2018; Kim et al. 2018; Martin et al. 2017). For example, European ancestry-derived polygenic scores have only 42% of the effect size in African ancestry samples (Duncan et al. 2019). Polygenic scores are also highly sensitive to inadequately controlled population stratification (Berg et al. 2019; Sohail et al. 2019; Morris et al. 2020). Even within a single ancestry group, the predictive accuracy of polygenic scores is dependent on age, sex and socio-economic status (Mostafavi et al. 2020)— this is not unexpected given the cultural variation that exists within a population (Muthukrishna et al. 2020; Muthukrishna and Henrich 2019). Similarly, the SNPs that contribute to the variance of a trait are different in different populations (Pemberton et al. 2018; Gurdasani et al. 2019; Akiyama et al. 2019; Rotimi et al. 2017) and it is difficult to disentangle the genetic, environmental and cultural contribution to differing polygenic scores between populations (Rosenberg et al. 2019). Recent projects have aimed to capture a greater degree of global human genetic diversity (e.g. Simons Genome Diversity Project Mallick et al. 2016; the exome analysis Lek et al. 2016; and the GenomeAsia project Wall et al. 2019), but we are far from accurately representing the genetic diversity of the global population.

## 3. BEHAVIORAL GENETIC PUZZLES IN LIGHT OF CULTURAL EVOLUTION

A dual inheritance and cultural evolutionary theoretical framework can help make sense of various puzzles in behavioral genetics. Here we discuss three: differences in heritability across socioeconomic levels, differences in heritability across development, and the Flynn effect.

### 3.1. Heritability across socioeconomic levels

The heritability of general intelligence is higher among affluent, high socioeconomic status (SES) households than poorer, low SES households in some societies (sometimes referred to as the Scarr-Rowe effect; Scarr-Salapatek 1971; Rowe, Jacobson, and Van den Oord 1999), but the correlation is mixed in other societies (Nisbett et al. 2012; Hanscombe et al. 2012; van der Sluis et al. 2008; Turkheimer et al. 2003; Giangrande et al. 2019; Platt et al. 2019). A meta-analysis (Tucker-Drob and Bates 2015) found the effect in a subset of US samples, but not in samples from Europe and Australia. Pooling the US studies, the authors found an effect size that corresponds to a heritability estimate of 0.61 at 2 standard deviations above the mean SES but only 0.26 at 2 standard deviations below the mean. In Europe and Australia, heritability is more uniform. The cause of this interaction is still debated.

Several researchers (e.g., Bates, Lewis, and Weiss 2013; Beam et al. 2015) have suggested that gene-environment correlation via phenotype-to-environment transmission, otherwise referred to as ‘reciprocal causation’ (Dickens and Flynn 2001; Bronfenbrenner and Ceci 1994; Scarr 1992), is the most likely explanation. By this explanation, those with genes well suited to a task can better nurture their skills in a wealthier environment than in a poorer environment. That is, initially small differences in genetic potential become gradually amplified over time due to the iterative matching of environments to abilities: an increase in expressed ability brings forth new environmental conditions that enable further growth along that dimension (Dickens and Flynn 2001; Bronfenbrenner and Ceci 1994; Scarr 1992). Such processes can increase genetic heritability, but through reciprocal shaping between genetic potential and environment, rather than through innately specified ability levels. The reasoning is that high-SES households are able to provide environments that do this more effectively and are thereby able to let genetic potential be more reliably associated with corresponding outcomes, lifting heritability as a result. While such reciprocal causation may indeed be occurring, reconciling this explanation with the findings from Europe and Australia seems more challenging.

A cultural evolution of genetic heritability explanation would instead suggest that heritability is a function of the variability in culture. In the United States, the differences between, for example, school and home environments among high SES households is small relative to differences between school and home environments among low SES households, where factors such as school lotteries can dramatically affect the cultural input. In contrast, the cultural environment is less unequal in Europe and Australia, where, for example, high quality schools are available across SES. Where these explanations make a different prediction is for poorer countries. The reciprocal causation explanation would predict low heritability in poorer countries. The cultural evolution of genetic heritability explanation would instead predict high heritability where there is equal access to similarly poor schools and households, but low heritability if inequality is high. That is, heritability is a direct function of variability in culture. More generally, we would predict a negative correlation between environmental variability and heritability (consistent with Davis et al. 2012).

### 3.2. Heritability across development

Cultural homogeneity may also vary across development. Because culture typically acts as a phenotypic homogenizer (see *Cultural compression problem*), we should be able to detect the influence of culture across development in the form of changes in genetic heritability. How we learn and who we learn from changes over the lifespan. One especially important transition is the shift from learning primarily from parents and other family members to learning from more distant models who are selected from a broader swath of society (Cavalli-Sforza and Feldman 1981). In the first of these two phases, there is less choice in what to learn, and much of the acquired knowledge is passed down through the same route as genetic information—from parent to child— by *vertical transmission*. In the second phase, the child is more independent, and has the opportunity to update what they have learned from a broader range of models, using learning strategies to decide whom to learn from, by *oblique transmission*. This expansion in learning models is essential for cumulative cultural evolution (Enquist et al. 2010).

This transition from vertical to oblique learning moves the child from the idiosyncrasies of their parents and household to the larger environment they now have in common with other adolescents and young adults. When the child is primarily relying on vertical transmission, the characteristics of their household plays a larger role in explaining variation in cultural input, in which case we should expect a high proportion of phenotypic variance to be explained by the shared (home) environment in twin studies. When the child switches to oblique learning, they now share more common influences with other children. This would be expected to reduce environmental variation across the population, and thereby increase heritability.

This reasoning implies that for phenotypic traits that are molded in real-time by the current shared environment instead of by the persisting effects of earlier parental influence, heritability should increase at this later life stage. Indeed, this is precisely what Hatemi et al. (2009) find in the case of political orientation, or where one lies on a progressive–conservative spectrum, measured in a US sample by questionnaire (Figure 4). Monozygotic (MZ) and dizygotic (DZ) twin pairs are both equally similar from middle childhood up to early adulthood, although the degree of twin similarity increases over time for both. Right around the age at which American children leave home, this pattern is broken, and the phenotypic correlations drop precipitously in DZ twins while remaining steady in MZ twins, and this discrepancy persists for the rest of the lifespan. The drop in DZ but not MZ correlation at this age suggests that the shared home environment exerts a convergent influence for both twin types early in life, but that once this influence is removed, genetic effects become unmasked and able to guide political attitudes independently from the shared environment. More phenotypic variance is explained by genes from this point onward, thus boosting heritability. In countries like Italy and Croatia, where the mean age of leaving the parental household is past 30 (European Statistical Office 2020), we would expect the developmental time course of heritability to reflect this later independence relative to American samples. Note that the present example has the same overall structure as the literacy example discussed earlier (Samuelsson et al. 2008), with heritability increasing as cultural influences from outside of the home environment kick in. Both examples indicate that heritability can be an index of shared life history and communal structure.

**Figure 4:**
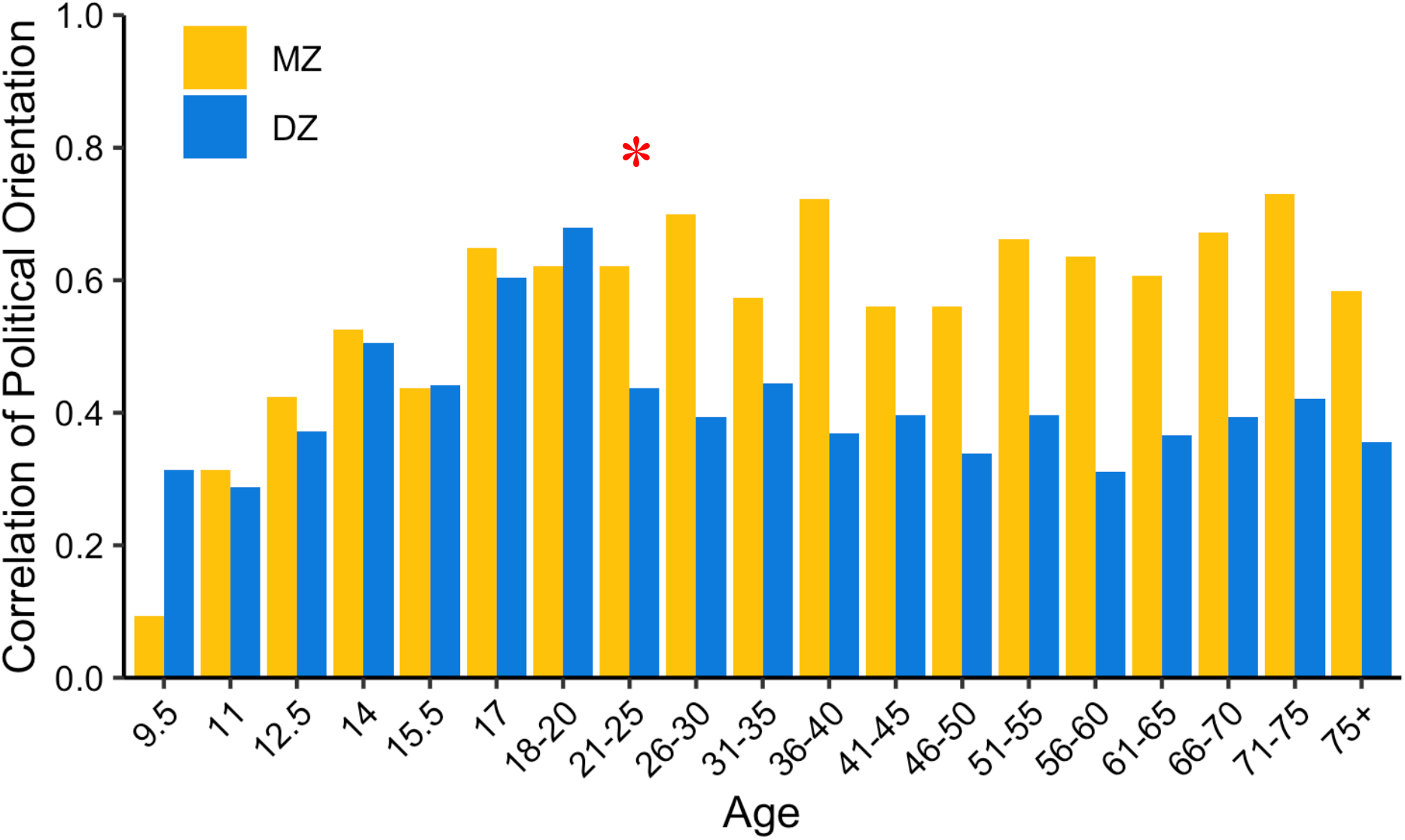
Twin concordances in political orientation. In middle to late childhood, within-twin correlations for reported political orientation are roughly the same between Monozygotic (MZ) and dizygotic (DZ) twin pairs in a US sample. In the early 20s, shortly after many US youth leave home for the first time to attend university, the correlation drops for the DZ twins but not for the MZ twins (identified with red asterisk). This shift corresponds to a sudden rise in heritability, as genetic similarity now predicts similarity in political orientation. When the effect of the home environment is weakened and replaced with more diverse cultural input, the effect of genes becomes unmasked and separates the phenotypic concordances between the two twin types. Horizontal axis indicates age, vertical axis indicates percentage twin concordances in political orientation. Figure reproduced from Hatemi et al. (2009).

Although the use of shared household environment to analyze twin data is a standard methodological convention, the household is in fact just one among many groupings of cultural organization that generate environmental convergence (Harris 1995). Households may be the most potent cultural grouping for some phenotypic traits, but other groupings may have significant impact as well, especially for specific kinds of traits. These may include schools, peer-groups, sports teams, religious communities; society-wide groupings such as mass media and popular culture; more diffuse groupings that are organized around particular sets of values such as political ideology or professional values; and possibly new kinds such as online communities. Although separating out the effect of household from the effect of genes is typically considered to be an explanatory goal, there may be further phenotypic variance that could be meaningfully explained if we were able to match phenotypes to relevant cultural groupings and gather data for both simultaneously. Twins share ‘common environments’ across multiple scales of social organization in this manner, but when phenotypic similarity is engendered by cultural groupings that extend beyond the household, the resulting correlations will usually be relegated to the broad category of ‘nonshared environment’ unless membership aligns with household structure (Plomin and Daniels 1987; Plomin, Asbury, and Dunn 2001). Although the nonshared environment is typically discussed as environmental exposure that is specific to the individual, it remains possible that there are multiple layers of communal structure embedded within this variance component. In societies such as the Israeli kibbutzim (Lieblich 2010), the Mosuo of southwest China (Ji et al. 2013), the Ache of eastern Paraguay, or the Hiwi of Venezeula (Hill and Hurtado 2009), children are raised communally and the groupings that are relevant for understanding genetic and environmental effects will be configured very differently from how the common environment is operationalized in the typical twin study.

General intelligence is another trait whose heritability is known to change over the course of development (Haworth et al. 2010; Briley and Tucker-Drob 2013; for a number of other traits see Bergen, Gardner, and Kendler 2007). This takes the form of a steady increase from childhood through adolescence all the way to early adulthood, after which it remains more or less steady over the lifespan. Although estimates vary, one meta-analysis (Haworth et al. 2010) put the heritability of general intelligence at 0.41 in childhood and 0.66 in adulthood. Explanations for this pattern typically invoke a combination of (1) gradual activation of relevant genes over the course of brain development and (2) active gene-environment correlation or ‘reciprocal causation’ (Bouchard 2013; Haworth et al. 2010; Plomin et al. 2016; Tucker-Drob, Briley, and Harden 2013). In contrast, a cultural evolutionary perspective would attribute the rise in the heritability of IQ to the developmental time course of cultural learning. We expect that in a society with different constraints on the development of cultural learning, the developmental trajectory of heritability would also differ.

A cultural evolutionary explanation of genetic heritability is consistent with this transition and would make an additional cross-cultural prediction—sharp changes in heritability will map onto sharp changes in an individual’s cultural environment (e.g. the start of school and university). These may shift due to policy changes, allowing for causal tests of this hypothesis. If, for example, children leave home later, then large increases in heritability should also be later.

### 3.3. The Flynn effect

The Flynn effect describes the rise in IQ test scores over time (Flynn 1984; 1987)—roughly 2 to 3 IQ points per decade on average around the world (Trahan et al. 2014; Pietschnig and Voracek 2015; Flynn 2009). The rate of increase differs between countries, being largest in countries that have recently started modernizing, and smallest in countries that had attained modernization by the beginning of the 20^th^ century (for review, see Nisbett et al. 2012). In some countries in Northern and Western Europe including Denmark, the Netherlands, and the United Kingdom, there is evidence that the Flynn effect has been slowing down and even reversing in recent decades (Dutton, van der Linden, and Lynn 2016). This negative Flynn effect is even less well understood than the positive Flynn effect. Bratsberg and Rogeberg (2018) find that in Norway, the negative Flynn Effect is found within families (between siblings), thereby making it unlikely to be explained by demographic changes or immigration, and instead supporting an environmental explanation.

There is no consensus regarding the cause of the Flynn effect, but given the recent and rapid increase, genetic explanations are unlikely. Common hypotheses include increases in test familiarity, improvements in education, sophistication of the technological and media environment, better nutrition, decreasing family size, and increased out-breeding or ‘hybrid vigor’ (Bratsberg and Rogeberg 2018; Clark, Lawlor-Savage, and Goghari 2016; Nisbett et al. 2012; Pietschnig and Voracek 2015; Trahan et al. 2014; Johnson 2006).

Flynn (2007) and Greenfield (1998; 2009) suggest that the effect is caused by a rapid worldwide increase of cultural practices, technologies, and environments that promote abstract cognitive processing as opposed to more traditional forms of concrete, pragmatic thinking. Some examples explored by these authors included urbanization, mass media, video games, education style, counterfactual thinking, and white-collar occupations. This account is mostly consistent with a cultural evolutionary explanation, which would suggest that intelligence is not just about hardware—genes, parasites, pathogens, pollution, and nutrition, but also software—the increasingly complex cultural package we acquire from our societies. By this account, not only is the idea of a culture-free IQ test implausible, but so too is the idea of culture-free IQ (for discussion, see Muthukrishna and Henrich 2016). Indeed, the largest Flynn effect can be seen on the supposedly culture-free Raven’s matrices (Nisbett et al. 2012; Flynn 2007), and on tests for fluid IQ rather than crystallized IQ (Pietschnig and Voracek 2015). When it comes to heritability, subtests of intelligence that are more culturally influenced are more heritable (Kan et al. 2013).

Beyond the diffusion of specific traits and abilities, a cultural evolutionary explanation also highlights how the Flynn effect is driven by the reorganization of cultural transmission pathways themselves. The introduction and improvement of formal schooling is one major instance of reorganization in cultural transmission that is also known to positively impact IQ (Ceci 1991; Davis 2014; Ritchie and Tucker-Drob 2018). Greenfield (1998) describes how IQ scores in some rural US towns in the early 20^th^ century increased rapidly at the same time as a number of coordinated changes in infrastructure, including better access to urban areas and new, high-quality road systems. Such enhancements in social connectivity directly translate into cultural connectivity, allowing for the influx and diffusion of psychological and behavioral traits that are considered valuable within the broader society. In much of the modern world, the kind of abstract information-processing ability measured by IQ tests is considered valuable, as it is useful in various white-collar professions that are typical of WEIRD societies. The Flynn effect therefore captures the progressive enhancements in cultural connectivity that have been occurring around the world because of improvements in various domains of infrastructure and technology including transportation, urbanization, education, and media. Global IQ rises in response to both the invention of relevant cultural traits and the enrichment of cultural transmission networks that carry those traits.

## 4. MODELING THE EFFECT OF CULTURAL EVOLUTION ON GENETIC HERITABILITY

### 4.1. Analytical model

Here we describe a simple mathematical model that captures the relationship between cultural evolution and heritability. Cultural evolution is a process in which some cultural variants spread through a population more prolifically than others. This spread can be partly due to intrinsic differences in the trait (e.g., steel axes are better than stone axes) and partly due to social learning strategies like the conformist bias, success bias, and prestige bias (for summary, see Kendal et al. 2018; Chudek, Muthukrishna, and Henrich 2015). Such strategies vary in their rules for selecting what to learn or whom to learn from, but they all lead to the disproportionate adoption of particular cultural variants over others, and thus to a reduction in the population-level variability of behaviors. Individual incremental improvement, individual learning (Rogers 1988; Legare and Nielsen 2015), cultural transmission error, and other sources of innovation (Muthukrishna and Henrich 2016) will continue to inject novel variants into the population, but the fact that a nearly unrestricted number of learners can inherit the behaviors and ideas of a few influential individuals makes it easy for cultural transmission to induce homogeneity. It is not only behavioral traits that become similar within a population through cultural transmission, but also environmental factors that shape behavioral traits, such as nutrition, sanitation, education, and media.

Heritability is defined as the proportion of phenotypic variance for some trait that is explained by genes. For broad-sense heritability, *H*^2^ = *V_G_*/*V_P_*, where *H*^2^ is heritability, *V_G_* is the variance in genotype and *V_P_* is the variance in phenotype. Because total phenotypic variance is made up of contributions from both genes and environment (*V_P_* = *V_G_* + *V_E_*), a reduction in the environmental contribution necessarily increases heritability. Heritability estimates are thus inflated for traits where environmental variance is reduced, such as through the homogenizing effect of cultural evolution. Behavioral geneticists partition phenotypic variance into genetic and environmental components, but here we further partition the environmental component into environmental variation unaffected by cultural evolution and environmental variation affected by cultural evolution. For convenience, we refer to the former as the ecological environmental variance and the latter as the cultural environmental variance, and represent this partition using the following notation:

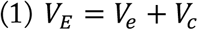

where *V_e_* and *V_c_* denote the variance in ecological and cultural environments, respectively. For simplicity, we model cultural environmental variation as a uniform continuous distribution that is bound by *k_min_*, the most unfavorable environmental state (for some given phenotype) within the experienced range of environments, and *k_max_*, the most favorable. We can use the variance of the continuous uniform distribution (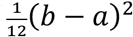, where *a* and *b* are the minimum and maximum values) to rewrite (1):

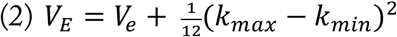

We can then substitute (2) into the standard formula for broad-sense heritability:

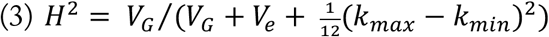

Heritability thus decreases when (*k_max_* − *k_min_*) is large and increases when it is small; the smaller the range of experienced states of the cultural environment, the more phenotypic variance there is left to be explained by genes and by consequence, the higher the heritability. The magnitude of this cultural effect depends upon (i) the ratio of *V_c_* to *V_e_*, which is the extent of cultural influence upon the phenotype-relevant aspects of the environment as a whole, as well as (ii) the ratio of genetic influence to total environmental influence (*V_G_* to *V_E_*). We illustrate the effect of each of these variance components on heritability in Figure 5.

**Figure 5:**
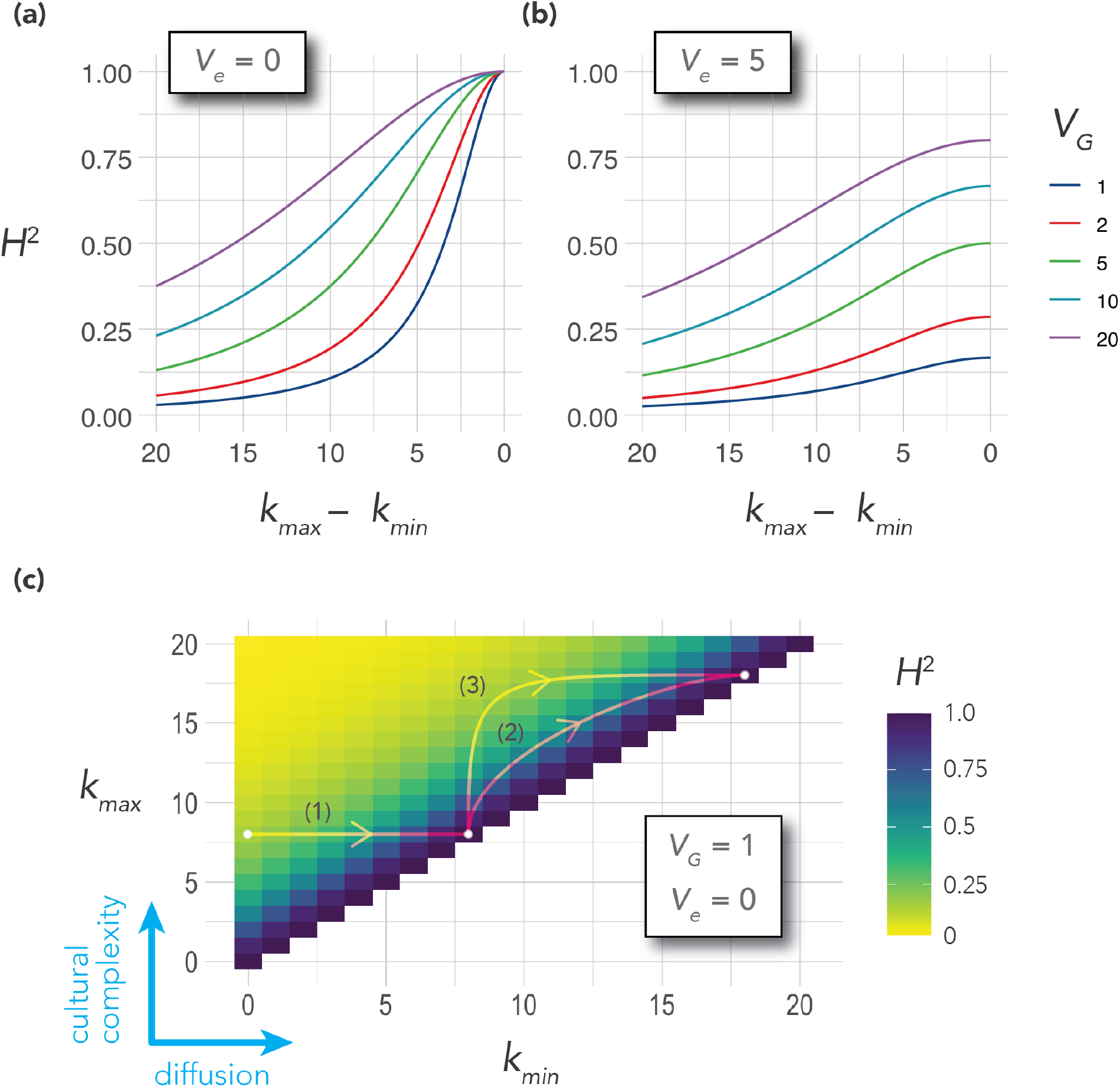
Visualizations of Equation 3. Heritability curves as a function of cultural range (***k_max_*** − ***k_min_***) and of the amount of genetic variance (***V_G_***). **(a)**Values are computed for ***V_e_*** = 0 (the environment is entirely explained by cultural factors) and **(b)**for ***V_e_*** = 5 (some of the environment is explained by non-cultural factors, such as climate). **(c)**An alternative visualization in which we look at the absolute values of ***k_max_*** and ***k_min_*** rather than just their difference, plotted for ***V_e_*** = 0 and ***V_G_*** = 1. An increase in ***k_max_*** expands environmental variation and implies increasing maximum cultural complexity, whereas an increase in ***k_min_*** reduces environmental variation and implies diffusion. Trajectory 1 represents a society’s transition from a more unequal cultural environment to a more equal cultural environment, but with no increase in cultural complexity. Trajectories 2 and 3 represent a simultaneous increase in cultural complexity and diffusion of the newly established complex traits, where a rising ***k_max_*** pulls ***k_min_*** upward but with varying lags: for trajectory 2 there is little lag between increase in the complexity of the culture and its spread, whereas for trajectory 3 there is considerable lag, with a longer period of relative cultural inequality. Genetic heritability decreases with rising cultural complexity and increases with cultural equality (diffusion). For example, continued innovation will reduce heritability up to the point at which the society is maximally unequal, and then increase heritability once more as the cultural innovations spread to the entire population—i.e., curves 2 and 3 are non-monotonic.

This model shows how heritability is not a fixed property of a trait, but rather a quantity that changes in relation to a shifting cultural environment. Although the model does not incorporate cultural dynamics as such, two aspects of cultural dynamics, increases in *cultural complexity* (Henrich 2004; Muthukrishna and Henrich 2016) and *diffusion* of those new innovations, are implicit in its formulation—new traits emerge and spread to fixation in the population. In the present model, we can think of increases in cultural complexity as pushing up *k_max_*, the most favorable cultural conditions in a society, and of diffusion as pushing up *k_min_* the most unfavorable cultural conditions in a society. As an example, imagine *k_max_* is the educational contribution of the best school in a society and *k_min_* is the educational contribution of the worst school in a society. In some societies, educational innovations diffuse quickly, whereas in others, there is more lag between the discovery of a new technology or pedagogical technique and its widespread adoption. Some societies are highly equal (*k_max_* − *k_min_* is small) and others are more unequal (*k_max_* − *k_min_* is large). The magnitude of the lag between increasing *k_max_* and *k_min_*, for example, how quickly educational innovations in the best schools diffuse to other schools, impacts environmental heterogeneity and thus heritability—we illustrate this effect as different trajectories of cultural change in Figure 5c.

### 4.2. Simulation model

#### 4.2.1. Mechanics

To capture the more fine-grained structure of the relationship between cultural evolution and heritability, we construct an agent-based model that simulates the underlying process. We then simulate the role of the behavioral geneticist by performing a “GWAS” on our constructed world and compare the recovered statistical associations to the underlying causal relationships. First, we create a population of agents who are allocated several ‘genetic’, ‘ecological’, and ‘cultural’ variables. Each of these variables, as well as the interactions among them, is assigned a fixed weight parameter that is drawn from a normal distribution with parameters *N*(0,1) (the cells of the coefficient matrix, Figure 6). That is, there are known main effects and interactions between genetic, environmental, and cultural variables on the phenotype.

**Figure 6:**
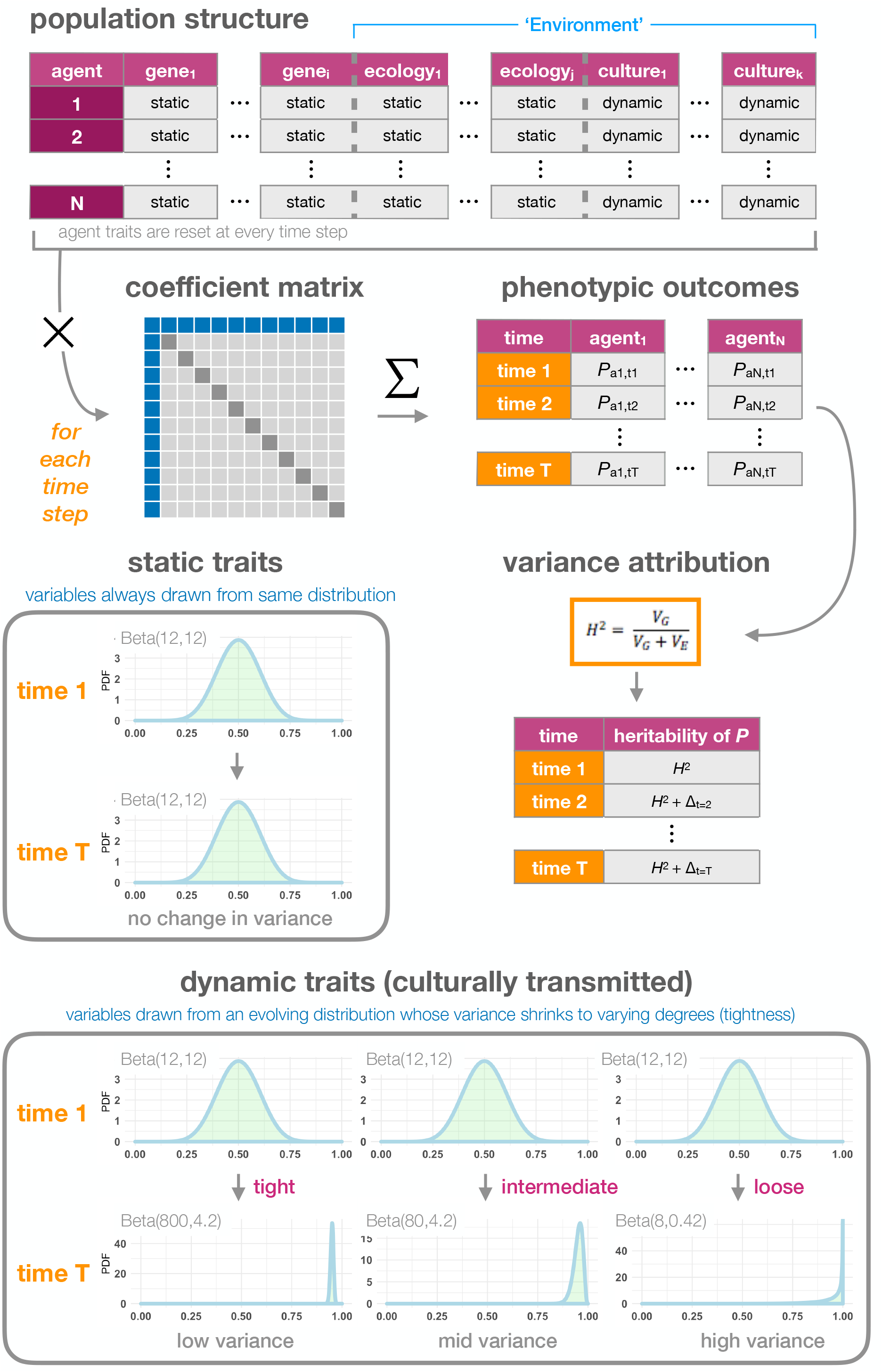
Set-up of the simulation model. A population of agents is allocated ‘genetic’ and ‘ecological’ variables that are drawn from a static distribution every time step, as well as ‘cultural’ variables that are drawn from an evolving distribution whose trajectory is dependent upon whether the society is ‘tight’, ‘intermediate’, or ‘loose’. Agent variables and all of their pair-wise interactions are multiplied by weights entered in a coefficient matrix and these weighted scores are summed to yield a single phenotype score for each agent. Heritability is computed every time step by running a linear regression on the main effects of the agent variables with respect to the phenotype score. Details are explained in the text.

Every generation, variables values are drawn for each agent and multiplied by the weight matrix to calculate phenotypic scores. Genetic and ecological variables are always drawn from a symmetrical beta distribution with fixed parameters Beta(12,12). Cultural variables are drawn from Beta distributions whose parameter values change smoothly over time modeling a process of cultural adaptation. Specifically, every cultural variable starts at Beta(12,12) which gives a mean of 0.5, but the distribution moves with each subsequent time step in a direction that maximizes the phenotypic outcome (on average). That is, if the weight matrix suggests a positive effect of a cultural trait, that trait moves toward fixation with the mean rising to 0.95. If the weight matrix suggests a negative effect of a cultural trait, the trait reduces, descending to 0.05 (Figure 6 show the increasing example).

The adaptive directionality models the outcome of a cultural evolutionary process as people selectively copy the majority or learn from common sources with a high payoff. We further model the *cultural tightness* of a society (Gelfand, Nishii, and Raver 2006; Gelfand et al. 2011) by changing the variance of the distribution (in addition to the mean change previously described). Tight societies, such as India and Singapore, enforce social norms and deter behavioral variation more than loose societies, such as Brazil and Australia, and thus tend to have relatively reduced variation in cultural traits. As such, in our model, variance shrinks the most for ‘tight’ societies and the least for ‘loose’ societies, with ‘intermediate’ societies in between.

We also model the role of a behavioral geneticist, who lacks knowledge of the causal weight matrix and must therefore infer the relationships that best explain the data. At each time step, we construct a linear model that regresses the phenotypic scores onto the genetic, ecological, and cultural variables using only main effects. GWAS typically only looks at main effects, although there has recently been an increasing focus on the detection of epistatic interactions (Niel et al. 2015). They also typically cannot measure ecological and cultural effects as we can in the model. Our behavioral geneticist compares the total phenotypic variance explained by genetic variables with the total variance explained by the combination of ecological and cultural variables (jointly, ‘environmental’ variables), and computes heritability using the standard formula. We thus obtain a record of the changing heritability of the phenotype over time, where this change is driven by the movement of the cultural distributions.

#### 4.2.2. Results

Although there is much variability across runs, as we would expect, heritability tends to rise across generations (Figure 7; Table 1) as a result of these cultural evolutionary dynamics. The magnitude of this rise is dependent upon society type, with the tight societies experiencing the most inflation and the loose ones experiencing the least. The tighter a society is, the more compressed its cultural variance, thereby reducing the amount of ‘environmental’ variance available to explain phenotypic variance and hence heritability. We also observe changes in the effect sizes of the predictor variables over time (example, Table 1): the coefficients captured by the regression model often move around in dramatic ways, strengthening or weakening over time, or at times reversing signs— much as different SNPs will have different effects depending on different conditions. The rank order of the contribution of the variables to the phenotype also fluctuates, as cultural evolution has differential impact across variables due to variation in the interaction weights.

**Figure 7:**
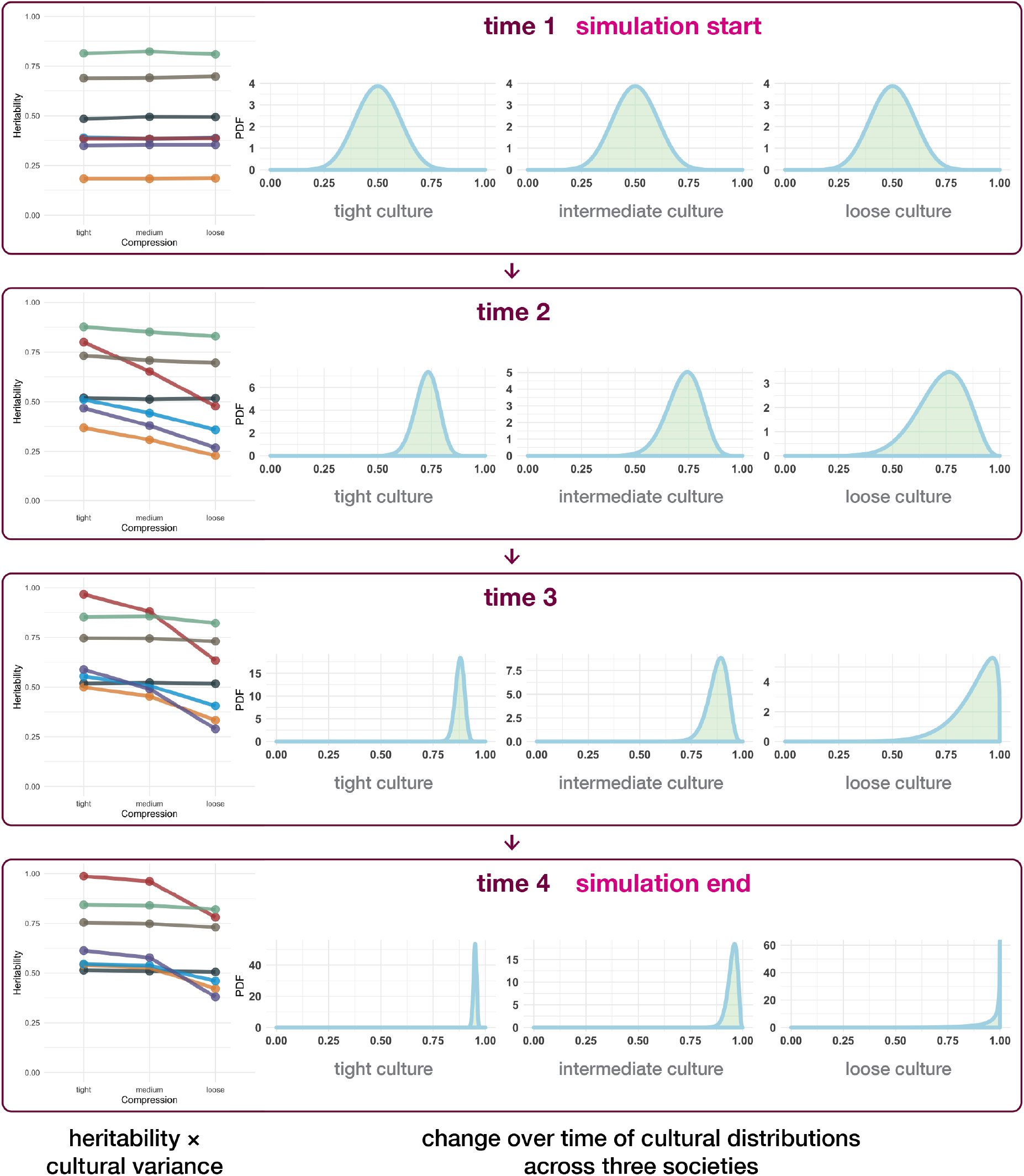
Simulation results for a set of example runs, displayed across 4 time steps (generations). Within each time step, the left-most panel plots heritability as a function of society type, with the left, middle, and right columns corresponding to ‘tight’, ‘intermediate’, and ‘loose’ societies, respectively. We randomly generated 7 configurations of the coefficient matrix, and for each configuration ran the simulation three times, varying only the society type for each. Same-color points connected by line segments indicate a common configuration. The three remaining panels within each time step display the shape of the Beta distributions from which the cultural variables are drawn, for tight, intermediate, and loose societies.

**Table 1:** Simulation output over one example run. Change in phenotypic variance explained by agent variables over 20 time steps (generations), in one example run of the simulation. Each row corresponds to a generation. The main block of ten columns represents genetic, ecological, and cultural variables, and the table entries are main effect coefficients for phenotype score regressed onto each variable, computed across all agents within a generation. Although only the cultural distributions are changing over time, the other variables get dragged along through their interactions with the cultural variables. Gene coefficients change here especially saliently for gene 3 which reverses sign and gene 4 which increases by a large amount. Toward the right are displayed the mean, variance, and heritability of the phenotypic score in each generation. Across all generations, the phenotypic score is calibrated to the distribution of scores in the first generation following the convention of IQ: a score of 100 is defined as the mean score of the first generation cohort, and a distance from the mean of 15 is defined as one standard deviation of the first generation’s scores (hence, variance is 15^2=225). Similarly to the Flynn effect, the mean phenotype score increases over time, in this case up 3.33 standard deviations from the first to last time steps, due to cultural adaptation. The variance shrinks rapidly and fluctuates over time. The heritability of the phenotypic score, or the proportion of its phenotypic variance that is explained by the four genetic variables, increases over time. This is due to the gradual reduction in the variability of the cultural variables, which is detectable in the shrinking magnitudes of their coefficients.

|  |  |  |  |  |  |  |  |  |  |  |  |  |  |
| --- | --- | --- | --- | --- | --- | --- | --- | --- | --- | --- | --- | --- | --- |
| 1 | -0.54 | -0.09 | -0.24 | 0.31 | -0.37 | -0.31 | -0.45 | 0.06 | -0.23 | 0.06 | 100 | 225 | 0.47 |
| 2 | -0.55 | -0.08 | -0.21 | 0.34 | -0.38 | -0.32 | -0.44 | 0.07 | -0.23 | 0.06 | 105 | 202 | 0.50 |
| 3 | -0.57 | -0.09 | -0.19 | 0.38 | -0.38 | -0.31 | -0.44 | 0.06 | -0.24 | 0.05 | 109 | 182 | 0.53 |
| 4 | -0.61 | -0.07 | -0.15 | 0.40 | -0.39 | -0.33 | -0.41 | 0.08 | -0.23 | 0.07 | 112 | 160 | 0.52 |
| 5 | -0.60 | -0.06 | -0.11 | 0.40 | -0.38 | -0.32 | -0.37 | 0.08 | -0.23 | 0.07 | 117 | 161 | 0.58 |
| 6 | -0.60 | -0.05 | -0.07 | 0.44 | -0.39 | -0.34 | -0.34 | 0.08 | -0.21 | 0.06 | 120 | 144 | 0.56 |
| 7 | -0.58 | -0.06 | -0.03 | 0.44 | -0.38 | -0.32 | -0.30 | 0.07 | -0.20 | 0.06 | 124 | 145 | 0.56 |
| 8 | -0.61 | -0.02 | 0.00 | 0.48 | -0.39 | -0.34 | -0.28 | 0.07 | -0.20 | 0.06 | 127 | 126 | 0.61 |
| 9 | -0.62 | -0.02 | 0.05 | 0.51 | -0.39 | -0.36 | -0.26 | 0.07 | -0.18 | 0.06 | 130 | 120 | 0.64 |
| 10 | -0.61 | -0.01 | 0.08 | 0.48 | -0.34 | -0.32 | -0.22 | 0.06 | -0.14 | 0.05 | 133 | 139 | 0.65 |
| 11 | -0.64 | -0.01 | 0.12 | 0.51 | -0.36 | -0.34 | -0.18 | 0.05 | -0.14 | 0.04 | 135 | 120 | 0.69 |
| 12 | -0.61 | 0.01 | 0.14 | 0.52 | -0.36 | -0.30 | -0.16 | 0.05 | -0.12 | 0.04 | 138 | 127 | 0.71 |
| 13 | -0.65 | 0.01 | 0.18 | 0.54 | -0.37 | -0.35 | -0.14 | 0.04 | -0.11 | 0.04 | 140 | 113 | 0.72 |
| 14 | -0.66 | 0.02 | 0.20 | 0.54 | -0.36 | -0.32 | -0.12 | 0.04 | -0.09 | 0.03 | 142 | 116 | 0.74 |
| 15 | -0.61 | 0.02 | 0.22 | 0.57 | -0.35 | -0.34 | -0.11 | 0.04 | -0.08 | 0.02 | 144 | 112 | 0.74 |
| 16 | -0.64 | 0.02 | 0.23 | 0.55 | -0.34 | -0.30 | -0.08 | 0.02 | -0.07 | 0.01 | 146 | 120 | 0.76 |
| 17 | -0.62 | 0.03 | 0.25 | 0.55 | -0.34 | -0.31 | -0.07 | 0.03 | -0.06 | 0.02 | 147 | 117 | 0.77 |
| 18 | -0.62 | 0.04 | 0.27 | 0.56 | -0.32 | -0.30 | -0.06 | 0.03 | -0.05 | 0.02 | 148 | 118 | 0.78 |
| 19 | -0.60 | 0.05 | 0.26 | 0.53 | -0.31 | -0.32 | -0.04 | 0.02 | -0.04 | 0.02 | 149 | 130 | 0.76 |
| 20 | -0.57 | 0.04 | 0.28 | 0.56 | -0.32 | -0.30 | -0.04 | 0.01 | -0.04 | 0.01 | 150 | 121 | 0.80 |
| Time | Gene 1 | Gene 2 | Gene 3 | Gene 4 | Ecological element 1 | Ecological element 2 | Cultural element 1 | Cultural element 2 | Cultural element 3 | Cultural element 4 | Phenotypic score mean | Phenotypic score variance | Phenotypic score heritability |

The mean phenotypic score increases over generations; recovering a Flynn effect (Table 1). In our simulation, the scores are pushed upward by the adaptively directed movement of the cultural distributions. The same mechanism also quasi-monotonically reduces the variance of the phenotypic scores over time. This is unlikely to be a characteristic of the empirical Flynn effect, whose sheer persistence over the span of a century (Pietschnig and Voracek 2015; although not everywhere, see Dutton, van der Linden, and Lynn 2016) indicates that it stems from continual waves of invention and diffusion, rather than a single diffusion wave as modeled in our simulation. Our model would therefore predict that during punctuated bursts of spreading innovation we should observe a pattern of increasing mean trait level coupled with decreasing variance.

The simulation described here is a minimal representation of how cultural evolution can influence heritability, even in the absence of genetic or ecological change. The magnitude of this influence is mediated by increasing homogeneity in the cultural environment, and also by the channeling of cultural change through environment–gene interactions that modulate the genetic effects of specific genes. We need to take these factors into account when assessing the effect of genes upon culturally malleable traits such as psychological and behavioral phenotypes. Genetic effects can be pushed around even for genes that are not under direct interaction with the cultural environment, because of fluctuation in relative standing due to cultural impacts on other genes. More generally, this model suggests a way to explain change in heritability over time, change in the effect size of specific genes over time, and change in phenotypic levels and variance over time (e.g., the Flynn effect), all within the scope of a single, tractable mechanism.

## 5. CULTURAL EVOLUTIONARY BEHAVIORAL GENETICS

Behavioral genetics offers a powerful empirical approach to understanding human behavior, but since the advent of whole-genome methods, its strategy appears to be predicated upon the notion that with enough data, the ground truth of human nature and nurture will be revealed. Data alone is not enough (Muthukrishna and Henrich 2019); the thrust of our theoretical case is that human psychology and behavior have a large cultural component that has been changing over history (Muthukrishna, Henrich, and Slingerland 2020; Nunn 2020; Henrich 2020; 2016; Chudek, Muthukrishna, and Henrich 2015; Laland 2018; Wilson 2019; Boyd and Richerson 1985; Boyd 2018). Most recently our psychology has been shaped by the advent of writing, numeracy, different types of agriculture, the Industrial Revolution, the Internet, and smart phones (Firth et al. 2019; Wilmer, Sherman, and Chein 2017; Ong 1982; Talhelm et al. 2014; Uskul, Kitayama, and Nisbett 2008; Domahs et al. 2010). As new adaptive traits emerge (Muthukrishna and Henrich 2016), initially those who possess these traits will have an advantage, as in the case of access to new food sources, better healthcare, more effective tools, or easier methods of learning. But eventually the traits will reach fixation in the population through the processes of cultural transmission (Henrich and Broesch 2011; Muthukrishna, Morgan, and Henrich 2016), at least until they are unseated by subsequent innovations (Kolodny, Creanza, and Feldman 2015; Muthukrishna and Henrich 2016). We predict that these cultural dynamics are reflected in heritability estimates. The more unequal the access to common culture, the lower the heritability, and by corollary, the more equal the access to common culture, the higher the heritability. When an innovation emerges, heritability may decrease initially, but will increase as the trait spreads through the population. Of course the rate and spread of innovations is ever faster today (Muthukrishna and Henrich 2016; Comin and Hobijn 2010).

As any geneticist knows, genetic heritability is a function of the variability in the environment, variability in genes, and variability in the phenotype. There is little to predict if the phenotype is homogenous, as in the number of fingers or kidneys. There is little to predict *with* if the environment or genes are homogenous. But what is factored into the environment includes not only the physical ecology, but also the cultural environment. While variance in genes and ecology may be relatively stable, the variance in the cultural environment is continually changing through the processes of cultural evolution. Therefore, genetic heritability estimates are highly affected by the degree of cultural homogeneity, which is itself a function of factors that affect cultural transmission such as sociality, transmission fidelity, tolerance for variation, population structure, and social network topology (Derex and Mesoudi in press; Muthukrishna and Henrich 2016; Henrich 2016). Under most empirical conditions, behavioral genetics underestimates the contribution of culture, including in estimates of heritability. We don't disagree with the findings in these fields or the data used, but instead argue that more nuance is required in how they are interpreted. Our dual inheritance demands that a genetic account of human psychology and behavior must also account for culture and cultural evolution.

### 5.1. Toward a dynamic model of environment

We are surrounded by the products of culture yet are generally unaware of the generative processes that bring such complex objects and conditions into existence. Cultural transmission spans broad networks of interconnected individuals, as well as deep time scales of inheritance. Each individual experiences just a snapshot, making the global mechanics opaque. Instead, each of us is left with an intuition that *our* world is largely *the* world, which perhaps explains why the WEIRD people problem is underappreciated even a decade after publication of Henrich et al. (2010) (Nielsen et al. 2017; Pollet and Saxton 2019; Tiokhin et al. 2019; Barrett 2020). From this limited vantage point, we evaluate questions such as the relative contributions of nature versus nurture. But our understanding of ‘nurture’ remains fundamentally anchored in our restricted experience of being enculturated into a particular environment, which leads us to implicitly see environmental features shared by members of our community as factors to be held constant, while our variables of interest—be they the absence of a parent, a childhood illness, birth into nobility, or a polygenic score—become matched to outcomes in our predictive models. Such models may be informal or formal, either encountered in community gossip (“children raised by single parents usually become…”) or in scientific journals (“growth mindset interventions predict…”; Sisk et al. 2018^5^).

Our need for causal explanations (Penn and Povinelli 2007; Gopnik et al. 2004) meets our tendency to essentialize people and groups, where genes offer a better essentialist vehicle than the environment (Heine 2017; Dar-Nimrod and Heine 2011).

The importance of limiting behavioral genetic findings to the reference population was famously argued for by Lewontin (1970), and remains a caveat for the analysis of genetic effects. But it is far less appreciated that the reason why a multitude of phenotypic factors can be successfully held constant (or controlled for) in the first place is, in large part, due to the convergent force of cultural learning. Lewontin (1970), in his counterargument to Jensen’s (1969) highly controversial article on the genetic determination of IQ, used inbred corn and a uniformly acting nutrient solution as his rhetorical props for explaining the environmental sensitivity of genetic effects. It is worth noting that domesticated crops experience a more homogenous environment not by accident, but as a product of human cumulative culture. Lewontin’s example is an illustration of how culture can generate at times extreme phenotypic convergence in significant features of our environment (or the environment of our domesticated flora and fauna).

We are all aware of gene–environment interactions (Hunter 2005; Moffitt, Caspi, and Rutter 2005; Lewontin 1970b), but we still tend to focus on what is predictive in our statistical models, which are constructed in a particular population and environment but whose apparent lessons are generalized beyond these contexts (e.g. the effects of an educational intervention). These models typically do not capture how the relevant environments are distributed within and between populations or how (or why) one type of environment transitions into another—‘environment’ is simply given as an exogenous variable. The cultural evolutionary approach forces us to explicitly recognize that human environments do not just happen to fall into place; they are rather the outcome of a dynamic, adaptive process that responds to both environmental and genetic factors. The literature on gene-environment interaction already recognizes genes and environments as non-orthogonal, but dependencies between the two are likely to be tighter and more prevalent than would be expected in a culture-free framework.

If we are to accommodate culture, ‘environment’ can no longer be treated as a static projection plane over which active elements (i.e., genes and G×E interactions) drop their shadows. Instead, both genes and environment—the latter animated by cultural dynamics—are in motion with respect to each other (as an example, see language–brain coevolution; Christiansen 1994; Christiansen and Chater 2008; Deacon 1997; as well as cultural niche construction; K. N. Laland, Odling-Smee, and Feldman 2001; Kevin N. Laland and O’Brien 2011). An environment can be used as a reference frame against which to judge the effect of genes, but this is done for pragmatic purposes and not because environments are intrinsically fixed. We might take our cue from James Gibson’s contribution to the study of vision (Gibson 1979a): the visual system works not by reconstructing its environment from the perspective of a fixed camera, but rather by geometrically transforming its surrounding surfaces through the movement of an animated perceiver^6^. Gibson recognized that environmental change is not noise, but rather the very medium through which the scientist obtains knowledge about visual function. Our argument presents an analogous approach to the study of genes.

### 5.2. Toward a new understanding of intelligence

The genetic underpinnings of intelligence have roots going back to the beginning of behavioral genetics (Galton 1869; 1874) and have been fiercely debated since at least Jensen (1969) and Lewontin (1970). The topic remains contentious, but a dual inheritance perspective cuts through some of this debate. Here we summarize key points.

General intelligence appears heritable—often measured at around 0.4 in toddlers and increasing up to 0.7 or 0.8 in adults (Bouchard 2009; Bergen, Gardner, and Kendler 2007). But as we have discussed, a high heritability score does not necessarily tell us whether a trait is primarily genetic; high heritability can also be an indicator of environmental homogeneity. Intelligence is a function of both our hardware (brain) and our software (culture) (Hutchins 1995; Vygotsky 1980), and the software has been changing far more and far more rapidly than has the hardware (Uchiyama and Muthukrishna in press). Genes certainly contribute to the size and organization of our brains— indeed, the Cultural Brain Hypothesis predicts a strong selection pressure for larger brains (Muthukrishna et al. 2018), still evident in the emergency interventions needed to successfully birth big heads (Lipschuetz et al. 2015). But those genes are explaining residual phenotypic variation only after accounting for environmental factors that also affect the “quality” of neural hardware, such as nutrition (Lynn 1990; Stoch et al. 1982), parasites (Wieringa et al. 2011), air pollution (Zhang, Chen, and Zhang 2018), and lead poisoning (Needleman and Gatsonis 1990; Wasserman et al. 1997). All are known to influence intelligence, but in societies that have been able to minimize variation on such factors, the environmental effect is also minimized. And it is not only such physical and physiological variables: changes and inequality in the cultural package delivered by schooling (Ceci 1991; Davis 2014; Ritchie and Tucker-Drob 2018) and our ever more complex entertainment media (Johnson 2006) also reduce the variation to be explained.

Recent, high-powered GWAS have found that genes associated with intelligence are expressed predominantly in the central nervous system (Sniekers et al. 2017; Savage et al. 2018; Davies et al. 2018), but these findings only explain the residual variation that remains after cumulative culture has reduced variation across many other variables—such as pathogens, parasites, and the nutritional environment—that would otherwise account for huge portions of variation on IQ test performance. The expression of “intelligence genes” may cluster inside the head, but this expression profile cannot be meaningfully evaluated without first considering the prior contributions of cumulative culture, which are invisible to standard methods within behavioral genetics.

As cultural and environmental factors become more homogenous, as they tend to do in developed countries, all that is left to explain the phenotypic variance are variables like differences in brain size or organization—the ‘causal locus problem’. It is unclear how strong the effect of brains is relative to everything that has been masked by culture, and due to the ‘cultural compression problem’, there is no simple, objective way to measure such a ratio. Investigating gene–IQ associations across maximally diverse cultural groups would at least be a step in the right direction—currently we suffer an even more acute ‘WEIRD gene problem’. The vast majority of GWAS of general intelligence conducted to date have sampled from individuals of European ancestry (Butcher et al. 2008; Kirkpatrick et al. 2014; Sniekers et al. 2017; Lee et al. 2018)^7^.

Intelligence may be highly heritable, but this is not an indication of its basis in the genome rather than the environment. The cultural environment amplifies heritability, and the degree of this amplification covaries with the extent to which a society has been able to reduce variation in physical, physiological, and informational factors that impact the phenotype. Developed countries, almost by definition, have been most successful in reducing this variation. High heritability of intelligence is therefore most likely to reflect the effect of the cultural environment in these societies.

## 6. CONCLUSION

> *Genetics is indeed in a peculiarly favoured condition in that Providence has shielded the geneticist from many of the difficulties of a reliably controlled comparison. The different genotypes possible from the same mating have been beautifully randomised by the meiotic process. A more perfect control of conditions is scarcely possible, than that of different genotypes appearing in the same litter*.(Fisher 1952)

In the above quote, Sir Ronald Fisher exalts the inferential purity that is afforded by the powerful pairing of sexual recombination with simultaneous multiple birth, which conveniently flattens environmental variation. But of course, this purity becomes progressively degraded with age, as environmental effects channel offspring through different developmental trajectories. Even among inbred, genetically identical mice who cohabit an experimentally controlled space, self-organizing trajectories of environmental experience result in clear differentiation in phenotypes like exploration, sociality, play behavior, and postnatal neurogenesis (Freund et al. 2013; 2015).

Humans trajectories differentiate so much more. We inhabit almost every ecosystem on Earth, not by speciating as many animals do, but through cultural adaptation, opening up different developmental pathways in different societies. But even within a single society, our massive specialization leads to high levels of differentiation. Our genetic variation explains some of this, but we are the least genetically diverse great ape—two groups of chimpanzees in central Africa are more genetically different from each other than two groups of humans plucked from Europe and Asia (Prado-Martinez et al. 2013). Most of our diversity is cultural rather than genetic (Bell, Richerson, and McElreath 2009; Muthukrishna et al. 2020); culture drives much of our within-species phenotypic variation. At best, genetic effects can only be specified within the ambit of a specific cultural context. And because cultures are also evolving over time, cultural contexts also require a timestamp. Heritability is not a property of a trait in itself, because in the absence of a reference culture it is necessarily unstable.

Cultural evolution yields cultural clusters. Within each cluster, environments are relatively homogenous (the degree of homogeneity also varies cross-culturally—tightness-looseness; Gelfand, Nishii, and Raver 2006; Uz 2015). This relative homogeneity that we find within societies is coupled with pronounced heterogeneity between societies (Bell, Richerson, and McElreath 2009; Richerson et al. 2016). Extrapolating genetic effects beyond a species is obviously mistaken, but so too is extrapolating a genetic effect beyond a society. But this is what researchers have been doing since Galton, and it is ingrained in both our methodology and our thinking, culminating in the recent triumphalist discourse surrounding behavioral genetics and GWAS. The movement toward more diverse genomic data ought to make some of these problems more obvious, just as more diverse psychological data made the problems of WEIRD psychology more obvious. But here too, data alone will not solve the problem (Muthukrishna et al. 2020; Muthukrishna and Henrich 2019). The question is not whether genes or culture contribute more to a behavioral trait, as behavioral traits can only be understood as emergent products of our dual inheritance, genetic and cultural. Nothing in behavioral genetics makes sense except in the light of cultural evolution.

## ACKNOWLEDGEMENTS

We would like to thank Veronika Plant for illustrating Figure 2.

## Footnotes

1 Research within this framework also falls under culture-gene coevolutionary theory and the extended evolutionary synthesis (Laland et al. 2015; Boyd and Richerson 1985; Cavalli-Sforza and Feldman 1981).

2 Australia’s Slip! Slop! Slap! campaign encourages practices to reduce UV radiation exposure: “slip on a shirt, slop on sunscreen and slap on a hat”. More recently, it has been followed by the SunSmart program, which expanded upon the original message to additionally include “seek shade or shelter, and slide on sunglasses”.

3 A similar phenomenon has been found in medicine (Gurven and Lieberman 2020).

4 This is an improvement from 2009 where 96% of GWAS participants were of European ancestry (Need and Goldstein 2009).

5 As an aside, that growth mindset might only replicate among low SES or at-risk students (if it replicates at all) fits with the general point that prediction is a function of variability. In this case, where there is a deficit, interventions may work, but where there is not, the potential gains are marginal or non-existent.

6 “The standard approach to vision begins with the eye fixed and exposed to a momentary pattern of stimuli…The ecological approach to visual perception works from the opposite end. It begins with the flowing array of the observer who walks from one vista to another, moves around an object of interest, and can approach it for scrutiny, thus extracting the invariants that underlie the changing perspective structure” (Gibson 1979, p.303).

7 Only 2 of the 27 studies of intelligence and 1 of 16 studies of self-reported educational attainment listed on GWAS Catalog include participants from other populations (accessed 21st April 2020). As well as European ancestry, five studies of intelligence and two studies of self-reported educational attainment also included samples with “NR”, or not reported ancestry.

